# Low Density Lipoprotein Receptor-Related Protein 1 (LRP1) as an auxiliary host factor for RNA viruses including SARS-CoV-2

**DOI:** 10.1101/2022.02.17.480904

**Authors:** Stephanie Devignot, Tim Wai Sha, Thomas R. Burkard, Patrick Schmerer, Astrid Hagelkruys, Ali Mirazimi, Ulrich Elling, Josef M. Penninger, Friedemann Weber

## Abstract

Viruses with an RNA genome are often the cause of zoonotic infections. In order to identify novel pro-viral host cell factors, we screened a haploid insertion-mutagenized mouse embryonic cell library for clones that rendered them resistant to the zoonotic Rift Valley fever virus (RVFV; family *Phleboviridae*, order *Bunyavirales*). This screen returned the Low Density Lipoprotein Receptor-Related protein 1 (LRP1, or CD91) as top hit, a 600 kDa plasma membrane protein known to be involved in a wide variety of cell activities. Inactivation of LRP1 expression in human cells reduced RVFV RNA levels already at the attachment and entry stages of infection. Moreover, the role of LRP1 in promoting RVFV infection was dependent on physiological levels of cholesterol and on endocytosis. In the highly LRP1-positive human cell line HuH-7, LRP1 also promoted the early infection stages of Sandfly fever Sicilian virus (SFSV; family *Phleboviridae*, order *Bunyavirales*), La Crosse virus (LACV; family *Peribunyaviridae*, order *Bunyavirales*), had a minor effect on RNA levels during the late infection stages by vesicular stomatitis virus (VSV; family *Rhabdoviridae*, order *Mononegavirales*), whereas infection by Encephalomyocarditis virus (EMCV, family *Picornaviridae*) was entirely LRP1-independent. Moreover, siRNA experoments in human Calu-3 cells demonstrated that also SARS-CoV-2 infection benefitted from LRP1. Thus, we identified LRP1 as a host factor that supports infection by a spectrum of RNA viruses.

## Introduction

Pandemics, epidemics, and zoonotic spillover infections are often caused by enveloped RNA viruses [1–4]. These pathogens contain an RNA genome of positive-sense or negative-sense polarity that is encapsidated by a viral nucleoprotein and surrounded by a lipid bilayer containing transmembrane glycoproteins. Being intracellular parasites with a comparatively small genome, viruses are exploiting cellular functions for basically every step of their replication cycle. For this, they are subjugating a multitude of host factors by interaction with specific viral proteins, as it is exemplified by the large virus-cell protein interactomes e.g. of SARS-coronavirus 2 (SARS-CoV-2) or influenza virus [5, 6]. Although RNA viruses are phylogenetically very diverse, it is conceivable that they may use overlapping sets of cellular factors.

Rift Valley fever virus (RVFV; family *Phleboviridae*, order *Bunyavirales*) is an emerging zoonotic negative-strand RNA virus [7] that is listed by the WHO among the pathogens posing the greatest public health risk [3]. Using RVFV as a model, we aimed to identify host cell factors supporting the viral replication cycle. A mutagenized cell library was iteratively screened for clones that acquired resistance to the highly cytopathogenic RVFV as an indicative that the affected host gene is essential for infection. As top ranking hit emerged the Low Density Lipoprotein Receptor-Related protein 1 (LRP1), a large plasma membrane receptor that can bind and internalize more than 40 different ligands [8–12]. In subsequent experiments we found that LRP-1 enhances the ability of RVFV to attach to the cell surface and enter the cytoplasm by endocytosis, and was also an auxiliary host factor for several other RNA viruses including the human pathogenic coronavirus SARS-CoV-2.

## Results

### Forward genetic screen for genes supporting RVFV infection

As RVFV is highly cytolytic, we devised a forward genetic screen that is based on the positive selection of cells deficient in pro-viral genes. A genome-wide library of knockout haploid mouse embryonic stem cells (mESCs), derived from parthenogenetic mouse embryos [13], was generated by mutagenizing with a retroviral genetrap (Fig. 1A) that disrupts genes in a revertible manner [14]. Altogether 5*10E8 cells were mutagenized by transduction with a genetrap retrovirus at an MOI of 0.02 resulting in approximately 10E7 independent mutations, selected by neomycin and expanded. Infection of the parental, non-mutagenized haploid mESCs with the attenuated RVFV strain MP-12 [15] was productive (Fig. S1A) and caused cytopathic effect (CPE) (Fig. S1B). Also after infection of the ~75 million genetrap library cells (7.5 cells/mutation) most of the mutagenized mESCs underwent CPE, but surviving and proliferating cell clones became enriched over 17 days under infection pressure. After repeated cycles of infection and selection (Figs. 1B and S2), all the surviving cell clones were collected, and the mutagenized genes then identified by an inverted PCR / restriction digest assay on genomic cell DNA, followed by Next Generation Sequencing (Fig. S3A and B, table S1). The top-ranking gene, found to be genetrap-inserted many times independently (Fig. S4), was Low Density Lipoprotein Receptor-Related protein 1 (*lrp1*). LRP1 is an approximately 600 kDa plasma membrane protein with a 515 kDa extracellular part [8–12]. For validation, we employed our genome-wide Haplobank library of more than 100,000 haploid mESC clones that contain a revertible genetrap with individual barcodes [14]. We took a number of independent Haplobank clones that carried a genetrap insertion in an intron of *lrp1* (table 1). Moreover, Cre-Lox inversion of the genetrap was performed, so each clone exists in a wt and a knockout version (sister clones; Fig. 2A). The mutant or wt mESC clones from the Haplobank and their respective reverted sister clones (each labelled either with GFP or mCherry, respectively; see table 1), were then mixed at a 3:7 ratio and subjected to a growth competition assay under RVFV MP-12 infection. Figure 2B shows that in most cases, infected cell clones exhibit a growth advantage when the *lrp1* gene is inactivated. The extent of the growth advantage was different for the different cell clone pairs, but showed a trend towards a certain infection resistance of the LRP1 knockout cells. For comparison, we employed Haplobank cell clones mutated in previously identified pro-viral host factors of RVFV, namely prenyltransferase alpha subunit repeat containing 1 (PTAR1; Fig. 2C) [16], and the E3 ubiquitin ligase FBXW11 (Fig. 2D) [17, 18]. Interestingly, in our mESC system the inactivation of these published host factors presented a much weaker survival benefit under selection pressure by RVFV than the deletion of LRP1. Thus, the growth competition experiments were in line with the initial results of the forward genetic screen and indicate that LRP1 may play a role in the life cycle of RVFV.

**Table 1.** **List of clones from the Haplobank collection** (mouse haploid embryonic stem cells, AN3-12)

| Gene | Clone # | Ref. Haplobank | Mutagen | Genetrap orientation | GFP | Cre-mCherry |
| --- | --- | --- | --- | --- | --- | --- |
| *Lrp1* | 4 | 00800IE06 | retroviral | antisense | wt | KO |
|  | 5 | 01128IH08 | transposon | sense | KO | wt |
|  | 10 | 00354IA04 | lentiviral | sense | KO | wt |
|  | 13 | 00561IB08 | retroviral | sense | KO | wt |
| *Ptar* | 15 | 01224IG10 | transposon | antisense | wt | KO |
| *Fbxw11* | 12 | 00371IH05 | lentiviral | sense | KO | wt |
|  | 16 | 01258IE05 | transposon | antisense | wt | KO |

**Figure 1.**
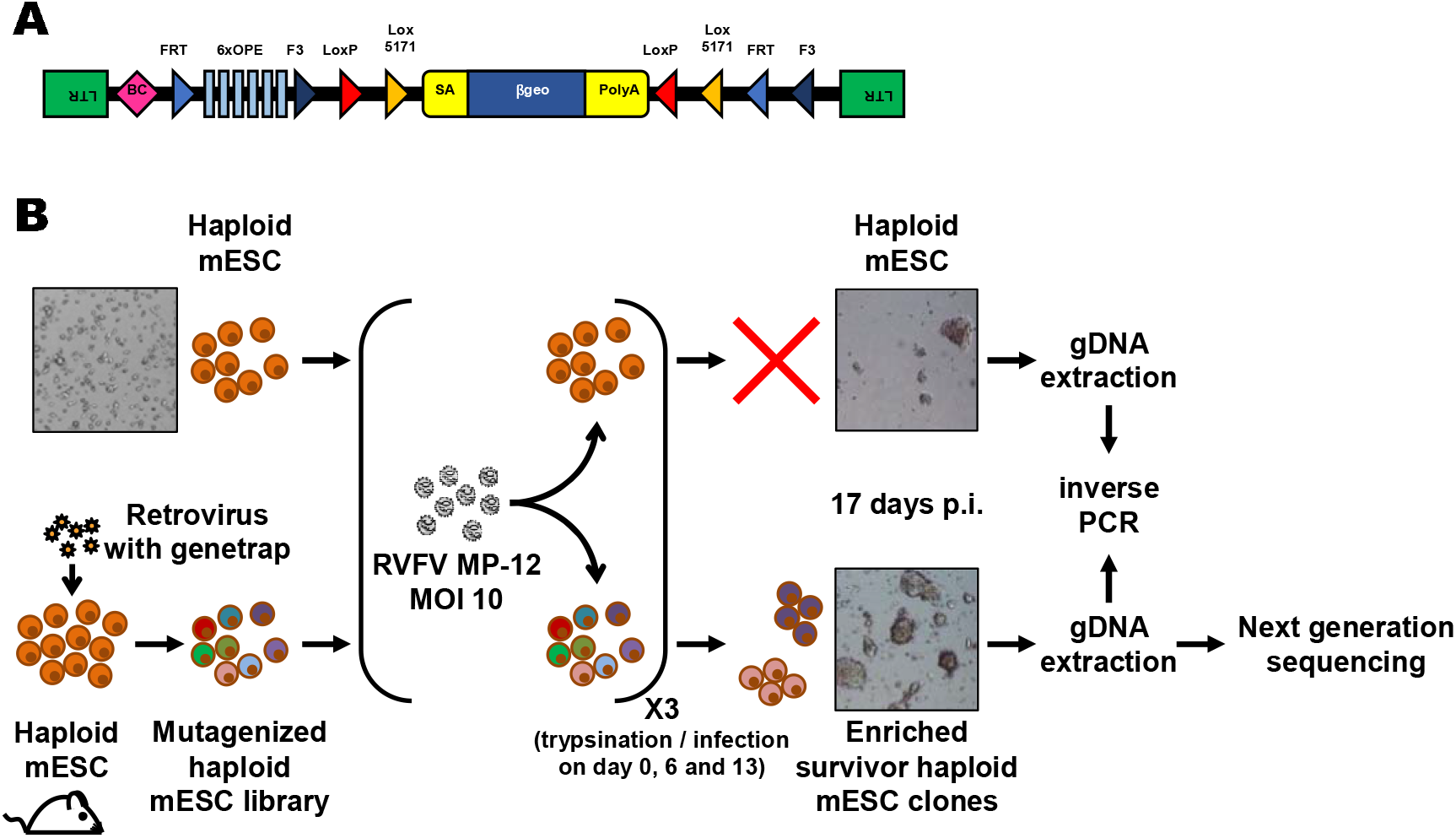
Forward genetics screen of the haploid mouse embryonic stem cell library for resistance to RVFV MP-12. (A) Schematic representation of the retroviral revertible genetrap used for mutagenesis. BC, barcode; OPE, Oct4 binding sites; SA, splicing acceptor site. (B) Experimental workflow for the RVFV MP-12 resistance screen using insertional mutagenesis. Bright-field microscopy images of the cells before infection and at the end of the screening process are shown as examples. mESC, mouse embryonic stem cells; p.i, post-infection; RVFV, Rift Valley fever virus.

**Figure 2.**
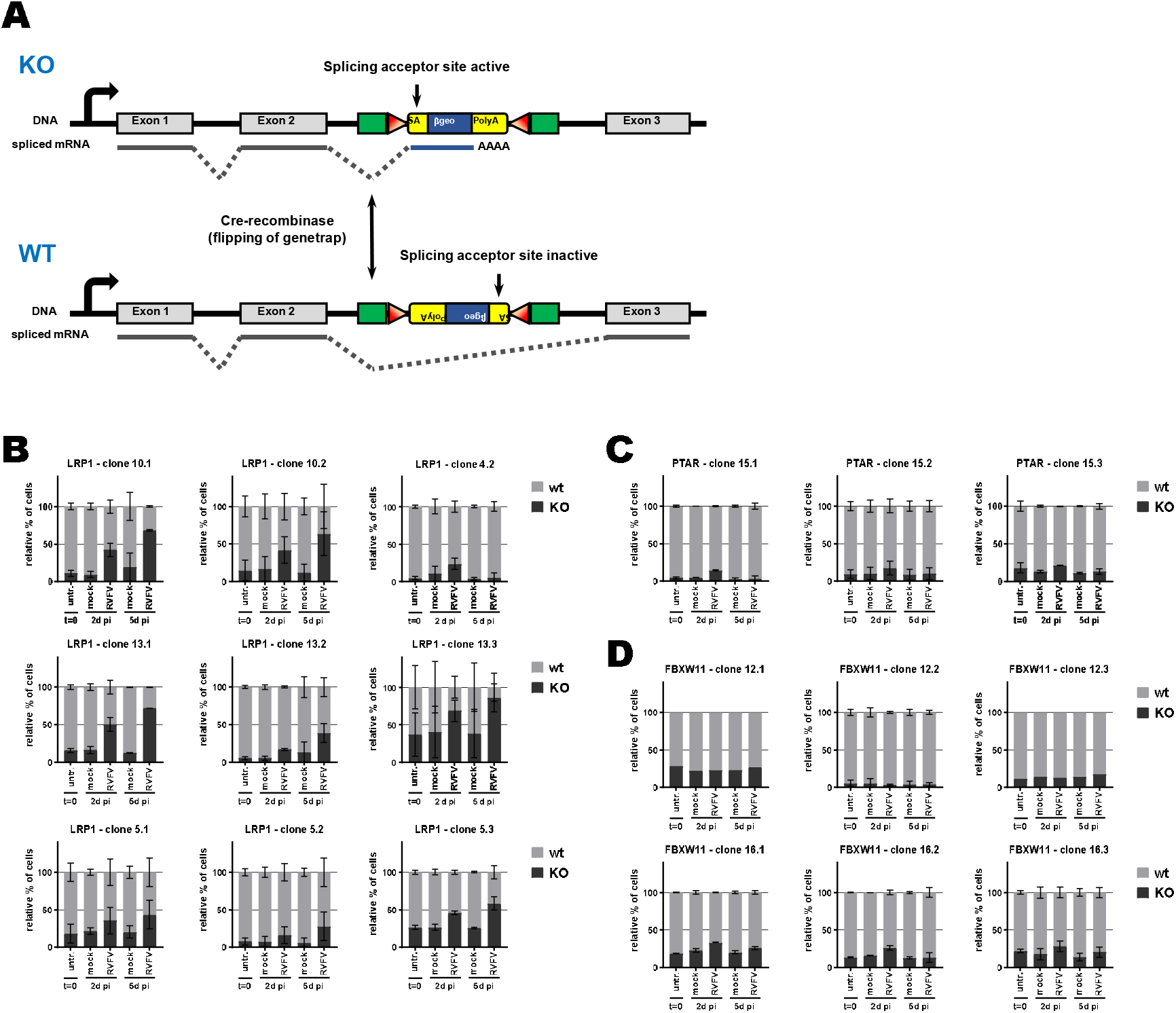
Growth competition assay in mouse embryonic stem cells. (A) Schematic representation of the genetrap system when inserted into an intron. When in sense orientation, the genetrap exposes a splicing acceptor site and will be inserted into the mature mRNA, leading to a knockout of the gene of interest. When in antisense orientation, the splicing acceptor site is inactive and the genetrap will be spliced out, leading to a wild-type expression of the gene of interest. Flipping of the genetrap orientation is possible by expressing the Cre-recombinase. (B, C, D) Growth competiton assay between sister clones bearing a genetrap into the gene of interest (B: LRP1, C: PTAR, D: FBXW11). Sister clones with wild-type phenotype are in grey and sister clones with knockout phenotype are in black (see table 1). Approximatively 30% of knockout cells were mixed with 70% of their wild-type sister clone, and infected with RVFV MP-12 at MOI 5. Ratio between both sister clones was followed by flow cytometry. n=2, except if event count was below 1,000, in which case the whole data set was removed (n=1)

Of note, our screen also returned two other top scoring genes, TBX3 and AIDA, but subsequent validations showed that unlike LRP1 they do not support RVFV replication (Fig. S5A and B).

### LRP1 impacts intracellular RNA levels of RVFV, but neither protein synthesis nor particle production

To follow up our findings from the mouse ESCs, we performed siRNA knockdown of LRP1 in human A549 cells and tested its effect on RVFV MP-12 replication using RT-qPCR [19]. siRNA-mediated downmodulation of LRP1 mRNA (Fig. S6A) resulted in an up to 50% reduction of RVFV gene expression at 5 h and 24 h p.i. (Fig. 3A), which is not accompanied by a concomitant increase in cell survival (Fig. S6B). Immunoblot analysis however revealed that A549 cells express comparatively little LRP1, whereas in human HuH-7 cells both the 515 kDa alpha chain as well as the 85 kDa beta chain gave strong signals (Fig. S6C). Therefore, we robustly downregulated LRP1 levels in the HuH-7 cells by introducing a CRISPR/Cas9 knockout (table S2 and Fig. S6C), and studied its phenotype with regard to RVFV replication. Also in the HuH-7 LRP1 knockout cells RT-qPCR analysis showed a suppression of viral gene expression by more than 50% already at 5 p.i. (Fig. 3B). However, levels of viral nucleocapsid (N) protein were unchanged between wt and LRP1 knockout cells (Fig. 3C and S5D). Moreover, when supernatants from the cells infected with MOI 0.01 were titrated, we did not observe any differences in virus yields. Thus, the phenotype of LRP1 deficiency under RVFV infection is measurable, but appears to be weaker in human cell lines than in mESCs, possibly because ESCs are in general more prone to virus infection due to their lack of an antiviral interferon system [20]. Altogether, we concluded from these data that LRP1 deficiency in human cell lines reduces RNA levels of RVFV, but that its absence seems to have no consequences for the production of viral N protein or progeny virus particles.

**Figure 3.**
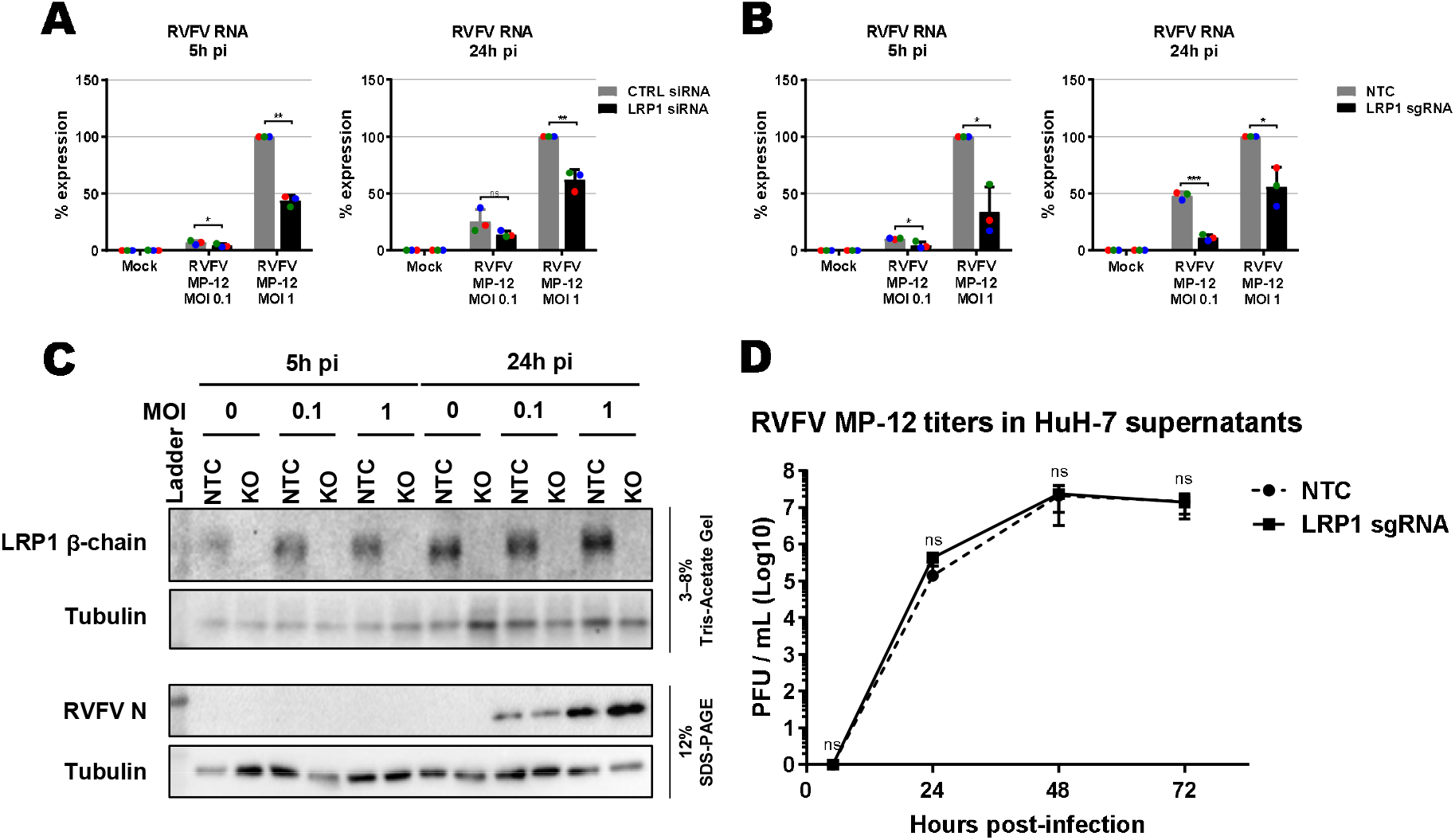
Influence of LRP1 downmodulation on RVFV MP-12. Virus RNA levels were measured in A549 knockdown for LRP1 (A) and HuH-7 knockout for LRP1 (B). Cells were infected in a synchronized manner with RVFV MP-12 at MOI 0.1 or MOI 1 as indicated, and RNA was extracted at 5 h and 24 h post-infection (p.i.). Two-step RT-qPCR was done to detect RVFV MP-12 RNA (L-segment) and the GAPDH reference gene. The RNA levels of RVFV L-segment in the control (CTRL) siRNA or no template control (NTC) CRISPR/Cas9 cells infected at MOI 1 were set to 100%. (C) LRP1 knockout HuH-7 cells were infected with RVFV MP-12 at MOI 0.1 or MOI 1, lysed at 5 h and 24 h p.i., and subjected to immunoblotting as indicated. A representative blot is shown. (D) HuH-7 CRISPR/Cas9 cells (NTC or LRP1 knockout) were infected with RVFV MP-12 at MOI 0.01, supernatants harvested at the indicated time points, and infectious virus measured by plaque assay. Statistics were done on 3 independent experiments, using a paired 1-tailed Student t-test (D: log-transformed data): *, p<0.05; **, p<0.01; ***, p<0.001; n.s., non-significant.

### LRP1 acts early in the RVFV infection cycle

LRP1 is a regulator of cholesterol homeostasis [21, 22], and cholesterol is important for virus infection [23]. We therefore investigated whether manipulation of cholesterol levels would influence the phenotype of LRP1 in RVFV infection. Cholesterol levels were either reduced by using Methyl-β-cyclodextrin (MBCD), or increased by enriching the incubation medium with cholesterol. As shown in figure 4A, MBCD treatment strongly decreased RVFV infection, as expected, and a difference between LRP1 positive and negative cells was not detectable any more. When the cells were given a surplus of cholesterol, infection was slightly reduced in LRP1 positive cells, but not in LRP1 negative cells, and also in this setting the difference between LRP1-deficient and -sufficient cells disappeared. Thus, the effect of LRP1 on RVFV infection appears to be dependent on physiological levels of cellular cholesterol.

**Figure 4.**
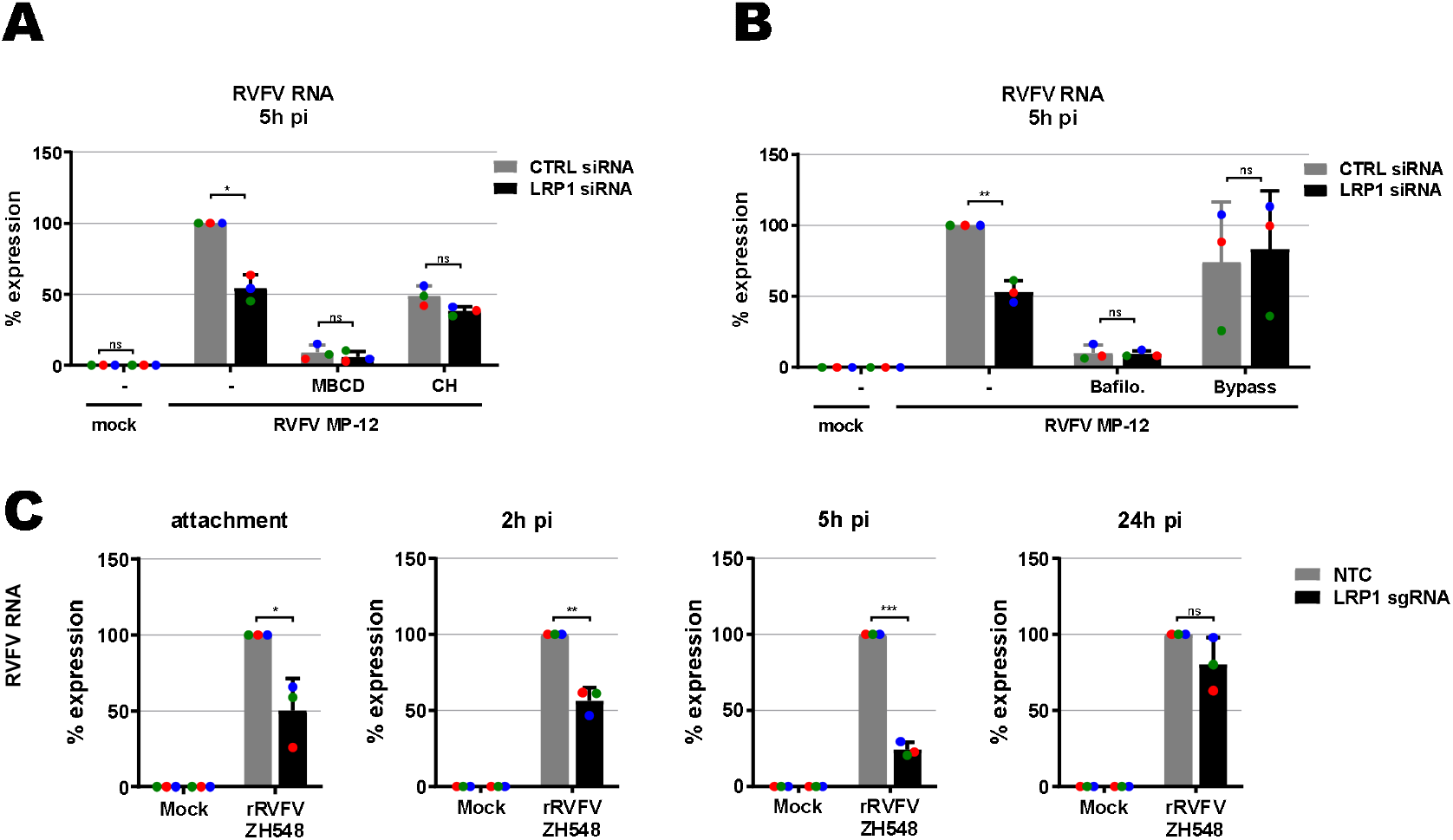
Mapping the LRP1-promoted infection steps. (A) Influence of cellular cholesterol levels. LRP1 siRNA-transfected A549 cells and their controls (see Fig. 3A) were pretreated for 1 h with Methyl-β-cyclodextrin (MBCD) to deplete cholesterol, or enriched with additional cholesterol (CH), before synchronized infection with RVFV MP-12 at an MOI of 1 for 5 h. (B) Role of endocytosis. Cells were pretreated for 1 h with bafilomycin A1 to block endosomal acidification, or incubated 3 min with acidic medium (pH 5.0) to force the fusion of viral particles at the cell surface (Bypass). (C) RVFV ZH548 RNA levels in LRP1 knockout cells over the course of infection. HuH-7 LRP1 knockout cells and HuH-7 NTC (no template control) cells were infected in a synchronized manner at MOI 1, washed 3 times, and further incubated in medium. Samples were collected after the 3 washes post-infection (attachment step), or at 2 h, 5 h or 24 h post-infection. Two-step RT-qPCR was done to detect viral RNA, as well as the GAPDH reference gene. The RNA levels in the infected NTC cells was set to 100%. Statistics were done on 3 independent experiments, using a paired 1-tailed Student t-test: *, p<0.05; **, p<0.01; ***, p<0.001; n.s., non-significant.

Like all bunyaviruses, RVFV enters the cells via endocytosis [24], so we wondered whether the LRP1 effect could be connected to this. Therefore, we either impeded endosomal acidification with bafilomycin A1, or bypassed endocytosis of RVFV particles altogether by acidification of the medium. The inhibition of acidification by bafilomycin A1 strongly impaired infection of the cells, whereas the endocytosis bypass did not substantially affect RVFV RNA levels, but wiped out the difference between the LRP1 positive and negative cells (Fig. 4B).

Our data suggest that the role of LRP1 in fostering RVFV infection is dependent on both cholesterol and on endocytosis early in infection. To determine the influence of LRP1 on all steps of virus replication, we infected HuH-7 wt and LRP1 knockout cells in a synchronized manner, and took cell-associated RNA samples at 0 h (attachment), 2 h (internalisation), 5 h (gene expression after entry into the cytoplasm) and 24 h (late phase of replication). For these analyses we engaged the wt RVFV strain ZH548 which unlike MP-12 is virulent for humans and animals [25]. As shown in figure 4C, LRP1 promotes already RVFV particle attachment and internalisation, and viral RNA levels in the mutant cells are lagging behind in the subsequent infection stages up to the 5 h p.i. time point. At the 24 h time point, however ZH548 RNA synthesis has recovered in the LRP1 knockout cells, different from what was observed to the attenuated mutant strain MP-12 (see Fig. 3A and B). Thus, in line with the results from the acidic bypass experiment, this indicates that LRP1 plays a role in the immediate-early steps of RVFV infection, namely cell attachment and entry. However, despite the differences that LRP1 made regarding viral RNA levels, there was no difference in virus yields (see Fig. 3D). It is therefore possible that particle assembly or budding are the rate-limiting steps in RVFV infection, and dominate over the weaker impact that LRP1 has on the immediate-early infection phase.

### LRP1 also supports other RNA viruses

The results from the knockout and time course experiments indicate a role of LRP1 as co-factor rather than as a main receptor of RVFV infection. We tested its importance for other RNA viruses, namely the closely related Sandfly fever Sicilian virus (SFSV; family *Phleboviridae*, order *Bunyavirales* ([26]), the more remotely related La Crosse bunyavirus (LACV; family *Peribunyaviridae*, order *Bunyavirales* [27]) and the non-related vesicular stomatitis (VSV; family *Rhabdoviridae*, order *Mononegavirales* [28]). Moreover, as these all are negative-strand RNA viruses, we additionally included the non-enveloped, positive-stranded Encephalomyocarditis virus (EMCV, family *Picornaviridae* [29]). As shown in figure 5 (A to E), LRP1 was also involved in particle attachment of the bunyaviruses SFSV and LACV. For these viruses the reduced infection in LRP1-deficient cells was more or less maintained throughout the replication cycle. By contrast, the rhabdovirus VSV and the picornavirus EMCV seem to attach to cells independently of LRP1. For EMCV, none of the replication stages was affected by a lack of LRP1, whereas VSV RNA levels were reduced at the 24 h p.i. time point of infection.

**Figure 5.**
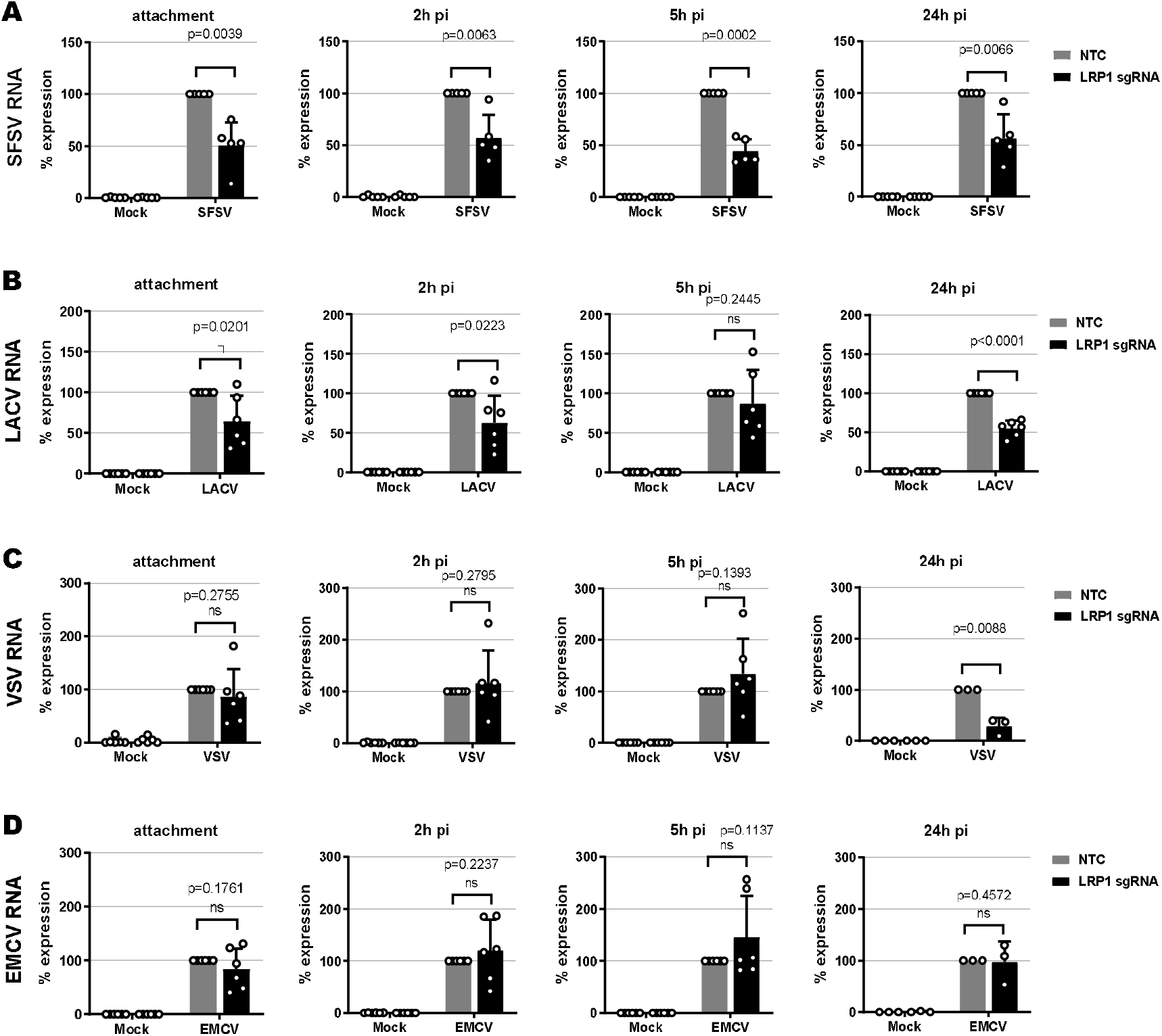
Virus RNA levels in LRP1 knockout cells over the course of infection. (A) Sandfly Sicilian virus (SFSV), (B) LaCrosse virus (LACV), (C) Vesicular stomatitis virus (VSV), (D) encephalomyocarditis virus (EMCV). HuH-7 LRP1 knockout cells and HuH-7 NTC (no template control) cells were infected with the various viruses at MOI 1, except for VSV that was used at MOI 0.1, washed 3 times and further incubated in medium. Samples were collected after the 3 washes post-infection (attachment step), or at 2 h, 5 h or 24 h post-infection. Two-step RT-qPCR was done to detect viral RNAs, as well as the GAPDH and 18S rRNA reference genes. The RNA levels in the infected NTC cells was set-up to 100%. Statistics were done on 6 independent experiments, using a paired 1-tailed Student t-tests: *, p<0.05; **, p<0.01; ***, p<0.001; n.s., non-significant.

Our comparative time course experiments in HuH-7 cells thus indicate that LRP1-dependency might be a common trait at least for bunyaviruses, but less so for the rhabdovirus VSV and not at all for the picornavirus EMCV. The fact that EMCV infection is unabated at all time points demonstrates that LRP1-deficient cells are in principle still able to support virus infection. For bunyaviruses, LRP1 is facilitating virus attachment, but the overall effect of its absence is comparatively low.

### SARS-CoV-2 infection of cells lacking LRP1

We also investigated the LRP-1 dependency of SARS-CoV-2, the causative agent of Coronavirus Disease 2019 (COVID-19) [30, 31]. For these experiments, we employed the human lung epithelial cell line Calu-3 because our HuH-7 CRISPR/Cas9 NTC and LRP1 knock-out cell clones exhibited differences in levels of the SARS-CoV-2 receptor ACE2 (Fig. S6E). When we performed siRNA knockdown in Calu-3 (Fig. S7A and B), LRP1 was found to be supporting RNA synthesis at 5 h p.i and 24 h p.i, but not at the earlier stages of the replication cycle (Fig. 6A). Moreover, a transient impact on viral N protein synthesis could be discerned in some of the replicates, but the effect was not statistically significant (Fig. 6B and C). In line with this, virus yields were comparable between wt and LRP1-deficient Calu-3 cells at all time points measured (Fig. 6D). The positive effect of LRP1 on SARS-CoV-2 infection therefore seems to be transient and limited to the viral RNA levels.

**Figure 6.**
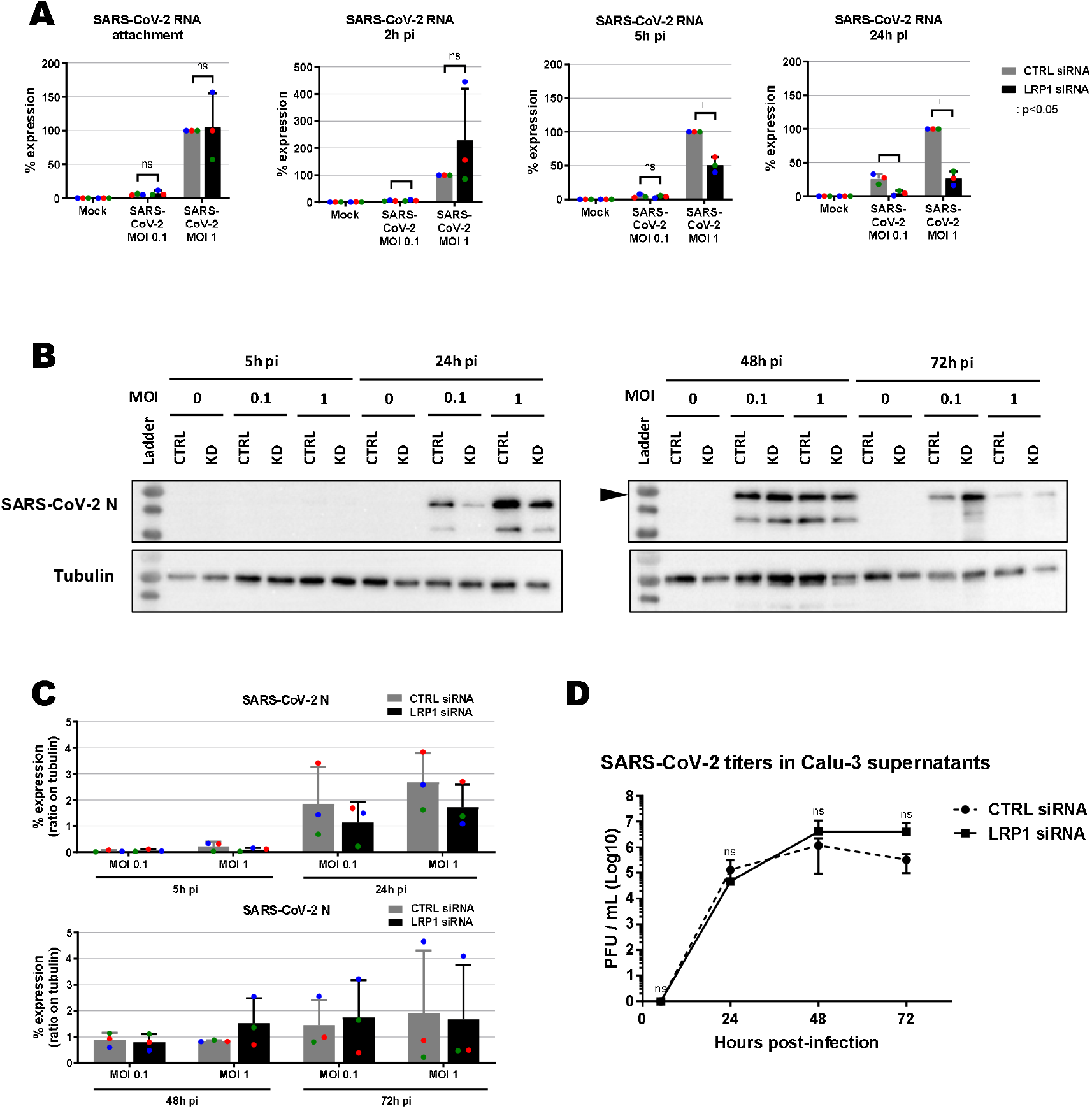
Effect of LRP1 knockdown on SARS-CoV-2 multiplication in Calu-3 lung cells. siRNA-transfected cells were infected at the two indicated MOIs, samples for viral RNA analysis (A), immunoblotting (B and C) and virus yields (MOI 0.01) (D) were taken at the different time points, and analyzed as described for figures 3 to 5. A representative blot is shown in (B). Quantifications of the N immunoblot signals relative to the tubulin signal are shown in (C). Statistics were done on 3 independent experiments, using a paired 1-tailed Student t-test (D: log-transformed data): *, p<0.05; **, p<0.01; ***, p<0.001; n.s., non-significant.

Thus, taken together, our forward genetic screen in haploid mESCs enabled us to identify the cellular protein LRP1 as promoting infection by RVFV and some other RNA viruses including SARS-CoV-2. Although the pro-viral effect of LRP1 was comparatively small, it was measurable, appears to be independent of the particular cell type, and mostly applied already in the attachment phase of the replication cycle. Thus, we conclude that LRP1 can act as an auxiliary host factor for enveloped RNA viruses.

## Discussion

LRP1 (or CD91) is a scavenger-type receptor involved in a wide variety of cell activities like migration, proliferation, differentiation, but also regulation of cholesterol homeostasis, inflammation, or clearance of plasma proteins from the blood stream [8–12]. It is important for the integrity of blood-brain barrier and can bind and internalize more than 40 different ligands (including apoptotic bodies or the Alzheimer disease-associated tau protein). Moreover, LRP1 modulates signalling pathways like e.g. those by JAK/STAT, ERK1/2 or TGF-ß [12, 32]. Our findings indicate that a series of RNA viruses are aided by LRP1 at their attachment and entry step, as might be expected from a membrane protein involved in constitutive endocytosis. Moreover, using RVFV as a model, we observe that cholesterol as well as endosomal acidification are involved in the positive influence of LRP1 on infection. In agreement with these findings, recent studies (published while our manuscript was in preparation) showed that LRP1 acts as a receptor for RVFV and the Oropouche orthobunyavirus (OROV) by binding to the viral envelope proteins and mediating their entry into the cytoplasm [33, 34]. Strikingly, LRP1 is a transporter that can overcome the blood-brain barrier [35]. Therefore, it may possibly be involved in the central nervous system tropism that is exhibited by viruses like RVFV, SFSV, LACV, or OROV [27, 36–38].

For two of our tested viruses – namely VSV and especially SARS-CoV-2 - LRP1 appeared to be only relevant for the later stages of infection. However, as the levels of viral RNA are low at the attachment and entry stages of infection, we cannot rule out that for these viruses the LRP1 effect is too small to be robustly detected at these early stages. Indeed, also Ganaie *et al*. investigated the LRP1 dependency of VSV and observed that the effect of LRP1 on VSV attachment and entry is negligible [33], but nonetheless a reduction of VSV spread in cell culture was detectable later in the infection cycle [34]. It is therefore conceivable that for some viruses hard-to-measure minor effects on attachment can amplify in a manner they become statistically robust only late in infection. On the other hand, coronaviruses like SARS-CoV-2 are known to intensively reorganize internal cell membranes in order to generate compartments that can serve as a safe space for transcription and replication of the genome RNA, and to assemble progeny particles [39–41]. As LPR1 is mostly cycling between the plasma membrane and the endosomes, it may transport factors critical for the RNA replication, e.g. lipids that are recruited for the formation of virally-induced intracellular membrane compartments. Hence, it appears possible that SARS-CoV-2 indeed engages LRP1 for late-stage replication, whereas for bunyaviruses it is attachment and entry. Future investigations should clarify whether and how SARS-CoV-2 is engaging LRP1 for its intracellular replication.

The results by us and others [33, 34] indicate that LRP1 is facilitating attachment and entry for a series of viruses. On the other hand we found that EMCV is not dependent on LRP1, and Schwarz et al. reported the same for the Zika flavivirus (ZIKV) [34]. Thus, LRP1 appears to be a broad, but not entirely general host factor for viruses. As EMCV is non-enveloped but ZIKV is enveloped, the dependency on LRP1 seems not to be due to the presence of a viral lipid envelope.

Our data as well as those of by Ganaie *et al*. and Schwarz *et al*.. [33, 34] clearly show that LRP1 is neither the only attachment factor for RVFV, nor for any of the other viruses tested. Indeed, heparan sulfates, C-type lectins like L-SIGN, and the intermediate filament vimentin were identified as auxiliary entry factors for RVFV, the LACV-related Schmallenberg virus, SARS-CoV-2 or many others [16, 42–49]. Interestingly, low-density lipoprotein receptor (LDLR), which belongs to the same protein family as LRP1 (“Low density lipoprotein receptor-related protein 1”) is the receptor for VSV [50], and has been proposed as an entry factor for flaviviruses and hepatitis B virus [51–53]. This may indicate that such endocytosis-active host factors are often exploited by viruses.

## Materials and Methods

### Cells and viruses

A549, BHK, HuH-7, Vero E6, Vero B4, as well as Calu-3 (kindly provided by Marcel Müller, Berlin, Germany) and LRP-1 knockout HuH-7 cells were grown in Dulbecco’s modified Eagle’s medium (DMEM) containing 10% foetal calf serum (FCS), 2◻mM glutamine, 100◻U/ml penicillin, and 100◻μg/ml streptomycin. Medium and supplements were purchased from Thermo Fisher Scientific. The mouse haploid embryonic stem cells AN3-12 were derived and sorted for their haploidy as described [13, 54]. The AN3-12 cells, the AN3-12 genetrap-mutated library, and the derived genetrap-mutated specific cell lines from the Haplobank [14] were grown in embryonic stem cell medium (ESCM) containing FCS and mouse Leukemia Inhibitory Factor (LIF) [14]. All AN3-12 cells were cultured in 10 cm cell culture dishes, passaged every second day by trypsinizing and re-seeding at 1:10, and the medium changed every other day.

Rift Valley fever virus (RVFV) strain MP-12, LaCrosse virus (LACV), Encephalomyocarditis virus strain FA (EMCV), and Vesicular Stomatitis Virus (VSV) were propagated in BHK cells. Recombinant RVFV (strain ZH548) and Severe Acute Respiratory Syndrome Coronavirus 2 (SARS-CoV-2 strain München-1.2/2020/984) were propagated in Vero E6 cells, and the Sandfly Sicilian virus (SFSV, strain Sabin) in Vero B4 cells. All virus stocks were confirmed to be mycoplasma-free. Infection experiments were done either under conditions of biosafety level 2 (BSL-2; EMCV, LACV, RVFV MP-12, SFSV, VSV) or level 3 (BSL-3; RVFV ZH548, SARS-CoV-2).

### Screening of haploid embryonic stem cell library with RVFV MP-12

We used a mouse haploid embryonic stem cell (mESC) barcoded-library (complexity: 9.7×10E6), mutagenized with the genetrap retrovirus JZ-BC frame 0, 1, 2 [14]. Thawed cells were grown overnight and seeded into seven 15 cm cell culture dishes at a density of 10×10E6 / dish. Four hours later, cells were washed with PBS and infected with RVFV MP-12 at an MOI of 10. As control, 3.9×10E6 genetrap-library cells and AN3-12 parental wild-type cells were seeded in 10 cm dishes, and infected by RVFV MP-12 at MOI 10, or incubated with the according mock supernatant. After 1 h at 37°C, infection was stopped by adding medium on top of the inoculum, and cells were further incubated at 37°C. Every 24 h during the whole screening process, medium was renewed and cell pictures were taken using bright field microscopy. At day 6 and day 13 post-infection (p.i.), surviving cells were trypsinized, seeded at same density for wild-type and genetrap-library cells, and re-infected 4 h later with RVFV MP-12 at MOI 10. At day 17 after the first infection, surviving cells were trypsinized and analysed for genetrap vector integration sites.

### Mapping of genomic genetrap vector integration sites

Experimental details of genomic DNA extraction, restriction digest, ring-ligation and inverse PCR with primers located in the genetrap (see Fig. S3A and B), as well as next generation sequencing of integration sites were described previously [13]. In short, the cell clones that were resistant to infection with RVFV MP-12 were trypsinized, washed in PBS and incubated overnight in gDNA Lysis buffer (GDLB: 10mM Tris-HCl pH 8.0, 5mM EDTA, 100mM NaCl, 1% SDS, freshly added 1 mg/mL Proteinase K). After treatment with 10μL RNase A (Qiagen) for 1 h at 37°C, genomic DNA was extracted with phenol/chloroform/isoamylalcohol and precipitated with isopropanol. Pellets were washed in 70% ethanol, dissolved in Tris/EDTA (TE) buffer, and the amount of DNA were measured using QuantiFluor dsDNA dye (Promega). An aliquot of 10 μg DNA was digested overnight at 37°C using *Nla*III and *Mse*I restriction enzymes (NEB), purified using Sera Mag Speedbeads (Thermo Scientific), and resuspended in TE buffer. After DNA ring ligation using T4 ligase (Roche) at 16°C overnight, DNAs were linearized by *Sbf*I (NEB) for 2 h at 37°C and further purified using Sera Mag Speedbeads. The integration site was amplified by inverse PCR (iPCR), using KenTaq polymerase (home-made), the primer DS, and one of the index primers (table S1): 3 min at 95°C, 36 cycles of [13 sec at 95°C, 25 sec at 61°C, 1 min 15 sec at 72°C], 5 min at 72°C, keep at 12°C. The iPCR products were visualised on agarose gel, and samples from the retro-library were purified with a Qiagen gel extraction kit.

Purified iPCR products (genetrap-library digested by either *Mse*I or *Nla*III) were quantified with a Nanodrop, and mixed 1:1 to be combined into one Next Generation Sequencing flowcell. Raw reads were trimmed to 50 nt and processed as in the NCBI Gene Expression Omnibus entry GSM2227065 [55]. In short, reads were aligned to the genome (mm10) with bowtie (v1.2.2) [56]. Insertions of disruptive and undisruptive regions of each gene are summed up (see Fig. S4). Binomial test of disruptive insertions against undisruptive regions and against disruptive insertions of a retrovirus input (GSM2227065), respectively, was performed for each gene. Genes were ranked by counts of disruptive insertions (DI) and were selected with a LOFscore <= 1e-20 (loss of function score). If the LOFscore equals 0, we remove genes with less than 10 DIs and without insertions in the background.

### Growth competition assay

Genetrap-mutagenized clones from the Haplobank collection (www.haplobank.at; [14]) were thawed, grown in 10 cm dishes, and split in 6 wells of a 24-well plate. Three of the wells were infected with a MLP-puro-GFP retrovirus, and 3 were infected with a MLP-mCherry-puro-Cre retrovirus (inducing flipping of the genetrap) [14]. At 24 h p.i., 1 μg/mL puromycin (Invitrogen) were added and 5 days later cells were split and aliquots were frozen.

For each gene of interest, cells of one GFP-labelled (original clone) and one mCherry-labelled (flipped sister-clone) version were mixed at a ratio of ~30% knockout cells to ~70% wild-type cells, respectively. At 4 h post-seeding, the mixed cells were washed with DMEM and either mock-infected or infected with RVFV MP-12 at MOI 5 for 1 h at 37°C. Medium (ESCM) was then added on top, and cells were further incubated at 37°C. Cells were trypsinized at 2 and 5 days pi, and either fixed in 4% PFA for flow cytometry analysis, or further grown after seeding in a new 24-well plate. The initial ratios between GFP and mCherry-labelled cells were confirmed, followed over-time by flow-cytometry (BD FACS LSR Fortessa, with HTS) and analysed with the FlowJo software. Only conditions with more than 1,000 acquired events were taken into account for final analysis.

### siRNA knockdown

A549 or Calu-3 cells were seeded in 6-well plates and reverse-transfected with LRP1 siRNAs Hs_LRP1_1 (SI00036190), Hs_LRP1_2 (SI00036197), Hs_LRP1_3 (SI00036204), and Hs_LRP1_5 (SI03109400) (FlexiTube GeneSolution, Qiagen) using Lipofectamine RNAiMAX reagent (Life Technologies), according to the supplier’s protocol. A second reverse-transfection was done 2 days later, and the cells seeded in 12-well plates before infection on the following day

### Generation of CRISPR-hSpCas9 knockout of HuH-7 cells

HuH-7 cells with a knockout in genes of interest were generated using the CRISPR-hSpCas9 strategy from the Zhang lab [57, 58]. For the *lrp1* gene, sgRNAs (see table S2) were designed using online tools (https://www.addgene.org/crispr/; http://www.e-crisp.org/E-CRISP/designcrispr.html). After cloning of the required plasmids, the lentiviruses expressing either the specific CRISPR-hSpCas9-sgRNAs or the no template control (NTC) CRISPR-hSpCas9 were generated, and then transduced in triplicates into HuH-7 cells. Clonal cell populations were isolated by limiting dilution, following the Addgene protocol (https://www.addgene.org/protocols/limiting-dilution/). Each single colony was further amplified, and screened by western blot. The control cells HuH-7 NTC (clone E5) and the HuH-7 LRP1 knockout cells (clone C8) were then used in further experiments.

### Reverse-transcription and quantitative PCR (RT-qPCR)

Total RNA was extracted from cell lysates using RNeasy (Qiagen) and an aliquot of 100 ng was reverse transcribed with the PrimeScript RT Reagent Kit with gDNA Eraser (Takara) and the included primer mix. An aliquot of 10 ng cDNA was used as template for amplifying sequences of human *GAPDH*, and *LRP1* and VSV with corresponding QuantiTect primers (Qiagen) or specific primers (table S3), respectively, and the SYBR premix Ex Taq (Tli RnaseH Plus) kit (Takara). *RRN18S* was amplified in a similar manner, but with 2 ng cDNA as template. RNA levels of EMCV, LACV, RVFV, SARS-CoV-2, and SFSV were detected using specific primers and Taqman probes (table S3), and the Premix Ex Taq (probe qPCR) kit (Takara). All PCRs were performed in a StepOne plus instrument (Applied Biosystems). The values obtained for each gene were normalized against GAPDH mRNA levels (or RRN18S mRNA levels in the case of the VSV RNA) using the threshold cycle (ΔΔ*C_T_*) method [59].

### Immunoblot analysis

Cells were washed with PBS and lysed in Tissue Protein Extraction Reagent (T-PER, Thermo Scientific) containing protease inhibitors (cOmplete tablets EasyPack, Roche), according to the supplier’s protocol. Samples for immunodetection of LRP1 were mixed with 4× sample buffer (143 mM Tris-HCl pH 6.8 (Acros), 4.7 % SDS (Roth), 28.6 % glycerol (Roth), 4.3 mM bromophenol blue (Roth)), while samples for detection of RVFV N or SARS-CoV-2 N additionally contained 20% beta-mercaptoethanol, and incubated for 10 min at 105°C.

For detection of LRP1, samples were run through Criterion XT Tris-acetate precast gels (3-8% gradient) (Biorad) in a XT tricine buffer (Biorad), for 1 h 20 min at 150V. Proteins were transferred on an EtOH-activated polyvinylidene fluoride (PVDF) membrane (Millipore) by wet blotting overnight at 40 mA and 4°C, using Tris-glycine transfer buffer (5.8 g/L Tris (Acros), 2 g/L glycine (Roth), 10% absolute EtOH (Roth)). The LRP1 α-chain (515 kDa) and β-chain (85 kDa) were detected using antibodies 8G1 (2μg/mL) (Merck-Millipore) and 5A6 (2μg/mL) (Merck-Millipore) respectively, and a horseradish peroxidase (HRP)-conjugated goat anti-mouse antibody (1:20,000) (Thermo Fisher).

For detection of RVFV N, SARS-CoV-2 N and ACE2 proteins, the samples were run through a home-made 12% SDS-PAGE gel for 1 h at 200V. Proteins were transferred on a MeOH-activated polyvinylidene fluoride (PVDF) membrane (Millipore) by semidry blotting for 1 h at 10V, using semidry blotting buffer (48 mM Tris (Acros), 39 mM glycine (Roth), 1.3 mM SDS (Roth), 20% MeOH (Roth)). The RVFV N was detected by the 10A7 antibody at 1.2ng/μL (kindly provided by A. Brun, INIA, Spain), and a horseradish peroxidase (HRP)-conjugated goat anti-mouse antibody (1:20,000) (Thermo Fisher). The SARS-CoV-2 N was detected by the anti-SARS-CoV nucleocapsid antibody (Biomol, ref 200-401-A50), and a HRP-conjugated goat anti-rabbit antibody (1:20,000) (Thermo Fisher). Human ACE2 was detected by the AF933-SP goat polyclonal antiserum (1 :1000) (R&D systems), and a HRP-conjugated donkey anti-goat antibody anti-goat antibody (1:20,000) (Thermo Fisher). Loading controls were performed by using an anti-tubulin antibody (1:4.000) (abcam, ref ab6046), and a horseradish peroxidase (HRP)-conjugated goat anti-rabbit antibody (1:20,000) (Thermo Fisher). Imunosignals were visualized using the SuperSignal West Femto kit (Pierce), and a ChemiDoc imaging system (Bio-Rad).

### Drug treatment

For assays involving cholesterol depletion or enrichment, cells were washed once with PBS, and pretreated 1 h at 37°C with either 5 mM Methyl-β-cyclodextrin (MBCD) (Sigma Aldrich, stock in ddH2O) in OptiMEM medium buffered with 25 mM HEPES (Sigma Aldrich, stock in ddH2O), or with 100 μg/mL cholesterol (Sigma Aldrich, stock in ddH2O) in OptiMEM, respectively. Cell monolayers were then washed 3 times in PBS, and infected as indicated above. After infection, cells were washed 3 times in PBS, and incubated in medium.

To manipulate entry of RVFV particles, cells were pretreated 1 h at 37°C with 20 nM bafilomycin A1 (Calbiochem, stock in DMSO) to block endocytosis. Cell monolayers were then washed 3 times in PBS, and infected as indicated above. After infection, cells were again washed 3 times in PBS, and incubated in medium supplemented with bafilomycin A1. To bypass the endocytosis step, cells were infected as indicated above, washed 3 times with PBS, and incubated 3 min at 37°C with pre-warmed acidic medium, pH 5.0 (adjusted with HCl 1M, Roth). The cells were then washed once with PBS, and further incubated at 37°C with normal medium. All samples were collected at 5 h p.i for further analysis.

### Infection time course analysis

Subconfluent cell monolayers were washed once with PBS, incubated 1 h at 4°C with the respective virus or a mock control and again washed 3 times with PBS. Samples for analysis of virus attachment were directly collected by incubating in RLT buffer (Qiagen) for 10 min at room temperature, resuspension, and storage at 4°C until RNA extraction. For analysis of the subsequent replication steps, medium was added and the cells were further incubated at 37°C. For the entry step, cells were washed once in PBS at 2 h p.i., incubated in trypsin-EDTA for 10 min (A549, Calu-3) or 3 min (HuH-7) to remove residual attached viral particles, and then washed 3 times in PBS with centrifugations (5 min at 10,000 g). Cell pellets were resuspended in RLT buffer and stored at 4°C. At 5 h and 24 h p.i., cells were washed once with PBS, incubated with RLT buffer for 10 min at room temperature, resuspended, and stored at 4°C.

### Virus titration

Supernatants of infected cells were collected and cleared by centrifugation at 400 g for 5 min. Serial dilutions were made to infect subconfluent Vero E6 monolayers, and infected cells were then incubated in medium containing 1.5 % Avicel for 3 days. Cells were washed twice in PBS and stained for 10 min with a cristal violet solution (0.75% crystal violet, 3.75% folmaldehyde, 20% ethanol, 1% methanol). Cells were then washed, and plaques were counted. Titers were determined as Plaque Forming Units (PFU) per ml.

### Statistical analysis

Statistical analyses performed are described in the figure legends.

## Acknowledgements

We are indebted to Christian Drosten and Marcel Müller for kindly providing the SARS-CoV-2 virus and Calu-3 cells, respectively, and to Andreas Leibbrandt, Ellen Wetzel, Manuela Kinzer and Nicole Schuller for helpful discussions and technical support. Work in the authors’ laboratories was funded by the Bundesministerium für Bildung und Forschung (Infect-ERA, grant “ESCential”) (FW, AM, JMP), the Swedish Research Council (VR; no. 2018-05766) (FW, AM, JMP), and the Pandemie Netzwerk Land Hessen (FW). Moreover, this project has received funding from the Innovative Medicines Initiative 2 Joint Undertaking under grant agreement No 101005026 (MAD-CoV-2) (FW, AM, JMP). This Joint Undertaking receives support from the European Union’s Horizon 2020 research and innovation programme and EFPIA.

## Author contributions

Conceptualization: SD, AM, UE, JMP, FW

Methodology: SD, AM, UE, JMP, FW

Investigation: SD, TWS, TB, PS

Visualization: SD, TWS, AM, UE, JMP, FW

Funding acquisition: AM, JMP, FW

Project administration: AH, JMP, FW

Supervision: UE, JMP, FW

Writing – original draft: FW

Writing – review & editing: SD, TB, AH, AM, UE, JMP, FW

## Data and materials availability

All data are available in the main text or the supplementary materials.

