## Supplemental Information for "Low Density Lipoprotein Receptor-Related Protein 1 (LRP1) as an auxiliary host factor for RNA viruses including SARS-CoV-2"

Figure S1

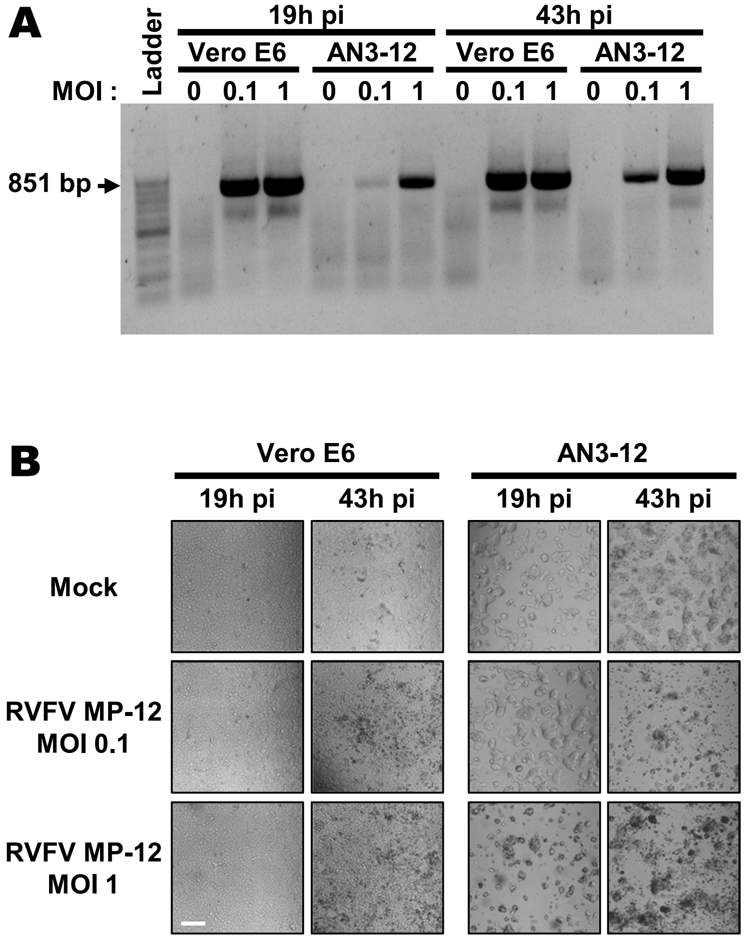


**Figure S1.** Replication of RVFV MP-12 in monkey kidney cells Vero E6, and mouse haploid embryonic stem cells AN3-12. (A) Two-step RT-PCR using primers located in RVFV S segment, flanking the NSs gene. (B) Bright-field microscopy images of infected cells. Scale bar: 0.5 mm.

Figure S2

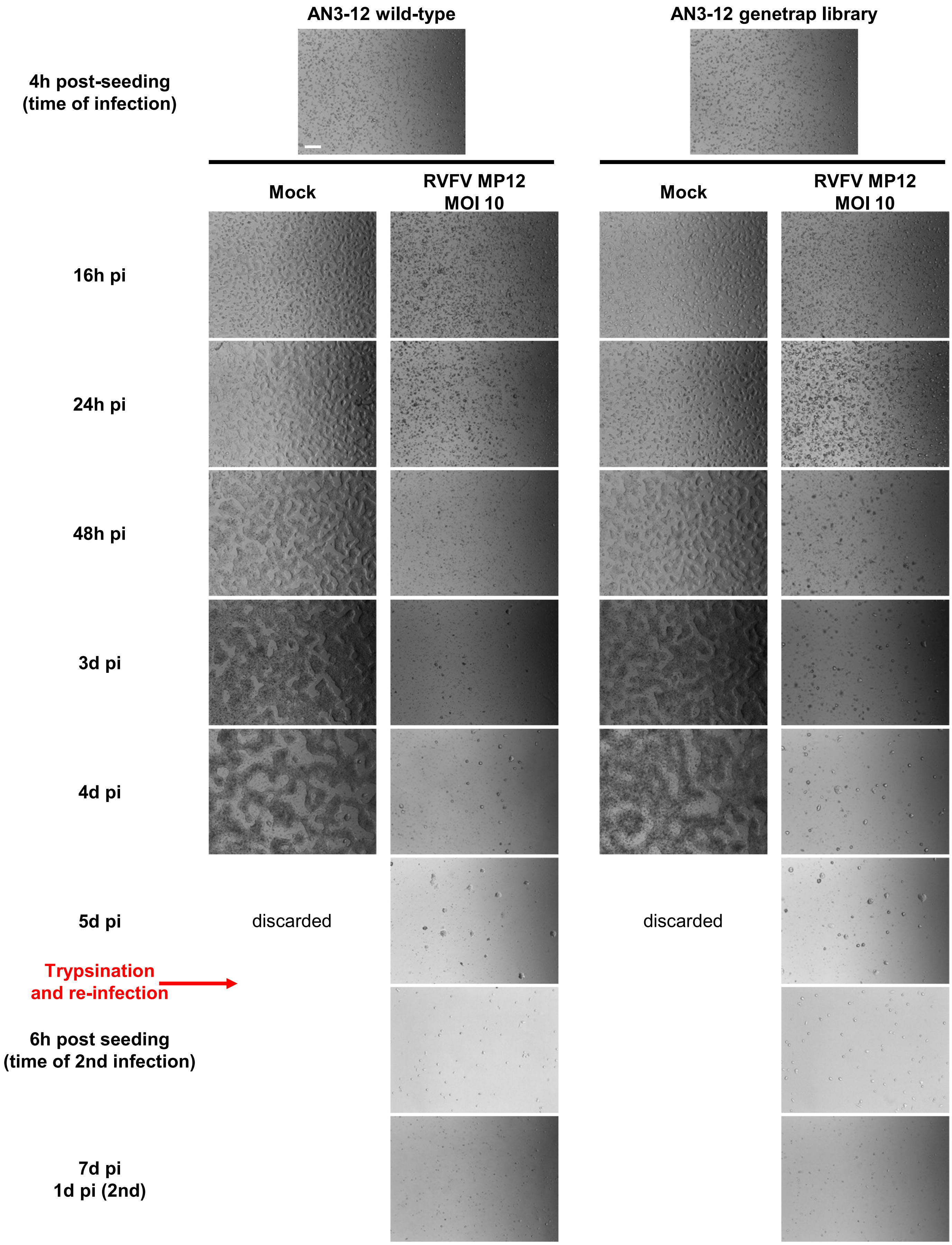
AN3-12 wild-type
AN3-12 genetrap library
4h post-seeding
(time of infection)
Mock
RVFV MP12
MOI 10
Mock
RVFV MP12
MOI 10
16h pi
24h pi
48h pi
3d pi
4d pi
5d pi
discarded
discarded
Trypsination
and re-infection
6h post seeding
(time of 2nd infection)
7d pi
1d pi (2nd)

Figure S2 (Cont.)

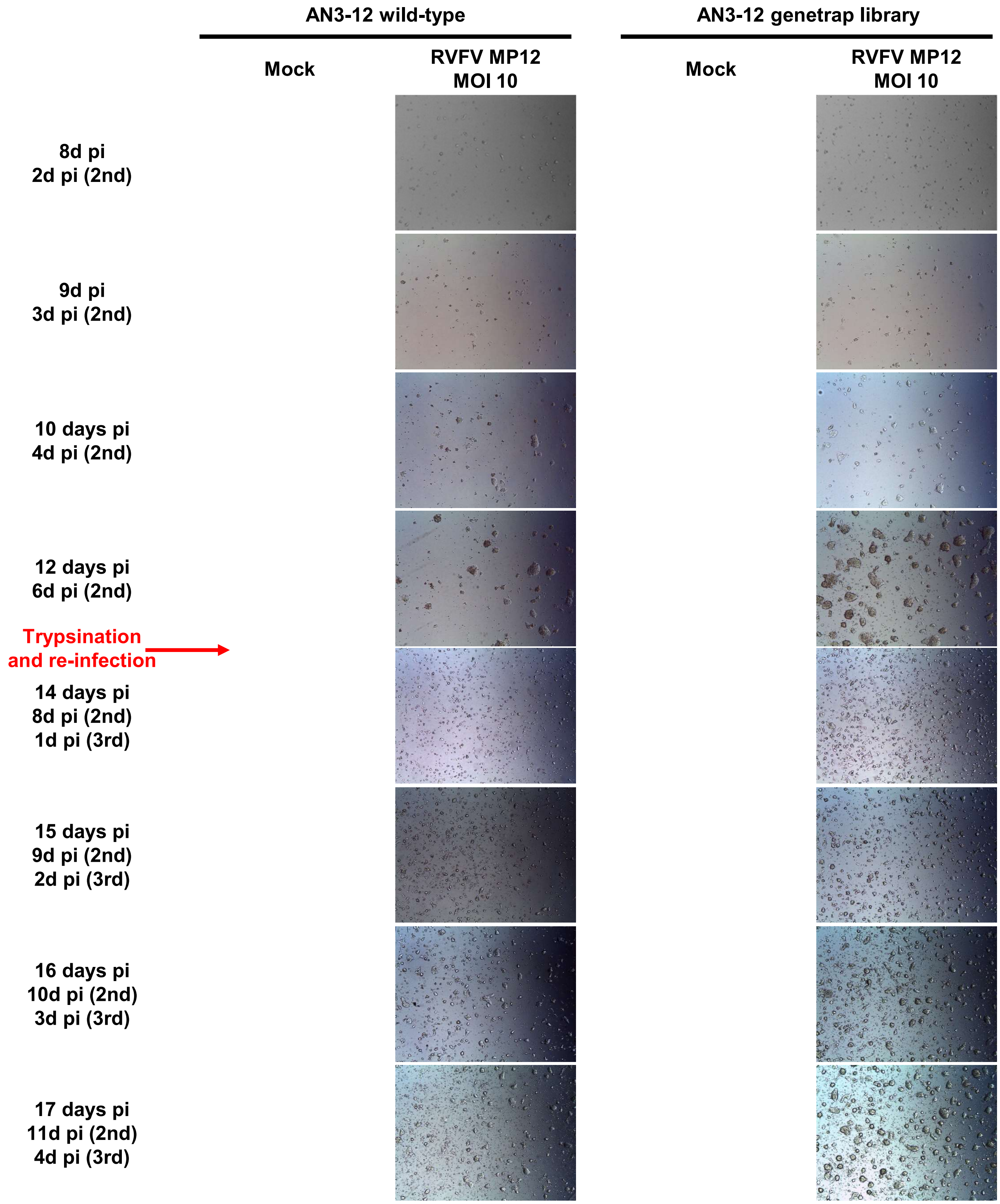


**Figure S2.** Bright-field microscopy of MP-12-infected mammalian embryonic stem cells, over the course of the forward genetic screen. Wild-type and retro-library AN3-12 cells were monitored every day over 17 days. Cells were trypsinized and reinfected at day 6 and day 13. A representative image is shown for each timepoint/condition. Scale bar (top left image): 0.5 mm.

Figure S3

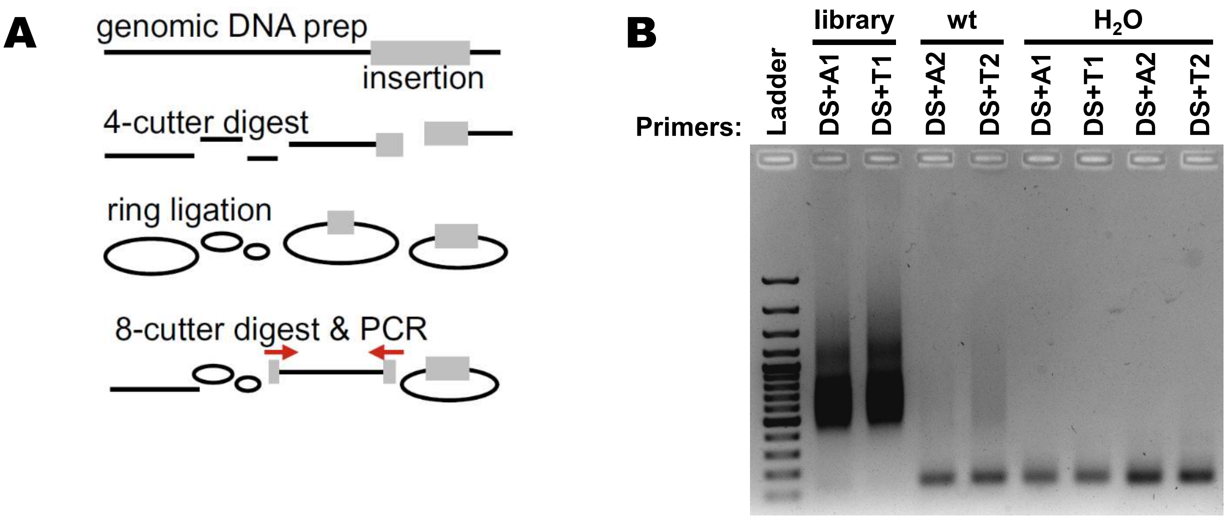


**Figure S3.** Inverted PCR (iPCR) on genomic DNA. (A) Schematic representation of the workflow process. The 4-cutter enzymes are MseI and NlaIII. The 8-cutter enzyme is SbfI. Primers are located in the genetrap and are designed to amplify the site of insertion. (B) Visualization of iPCR products in an agarose gel. Primers are described in Supplemental Table 1.

Figure S4

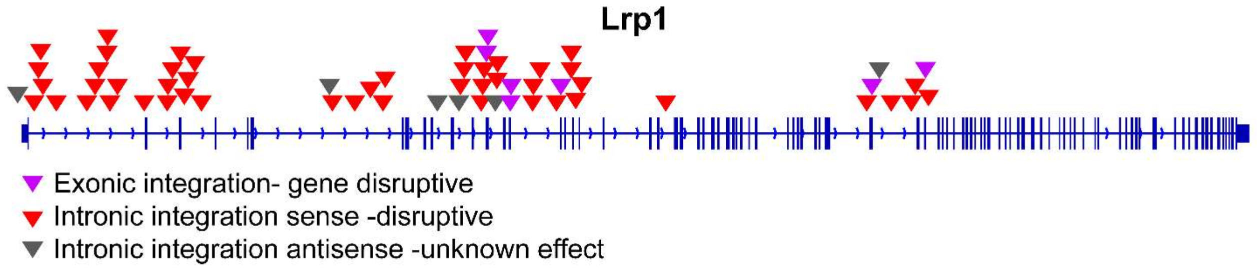


**Figure S4.** Next generation Sequencing analysis of NlaIII- and MseI-digested RVFV-infected AN3-12 library, showing the presence of the genetrap in the LRP-1 gene.

### Figure S5

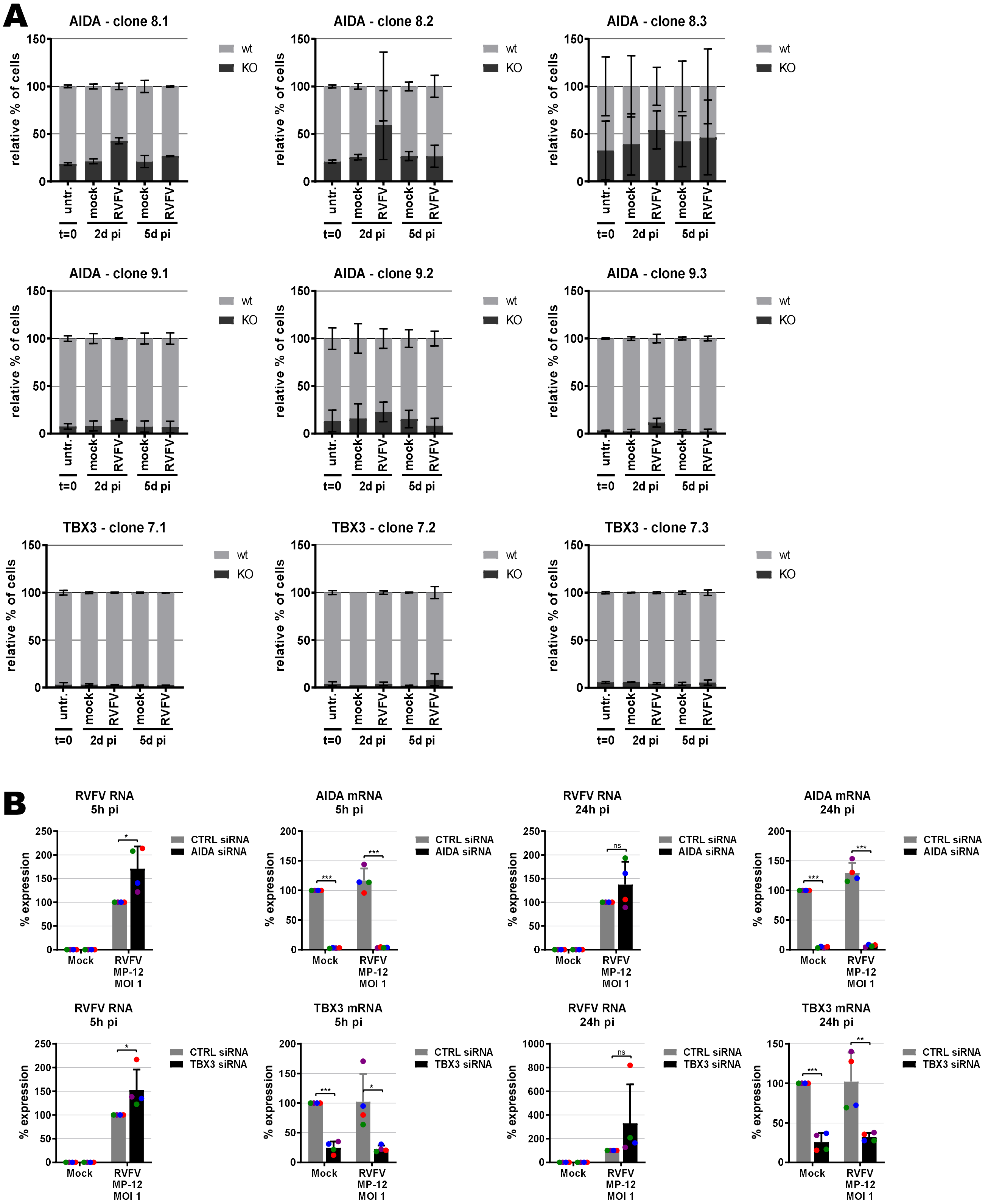


**Figure S5.** Validation of top scoring genes Tbx3 and Aida, using the same methods as described for LRP1 (see Fig.2). (A) Growth competition assay performed in triplicates, (B) siRNA knockdown and RT-PCR analysis for RVFV RNA (MOI 1, n=4). siRNAs against TBX3 (Hs_TBX3_3, Hs_TBX3_4, Hs_TBX3_5, Hs_TBX3_6) and AIDA (Hs_C1orf80_1, Hs_C1orf80_2, Hs_C1orf80_3, Hs_FLJ12806_4) were purchased from Qiagen, as were RT-qRNA primer pairs (Hs_AIDA_1_SG, Hs_TBX3_1_SG).

Figure S6

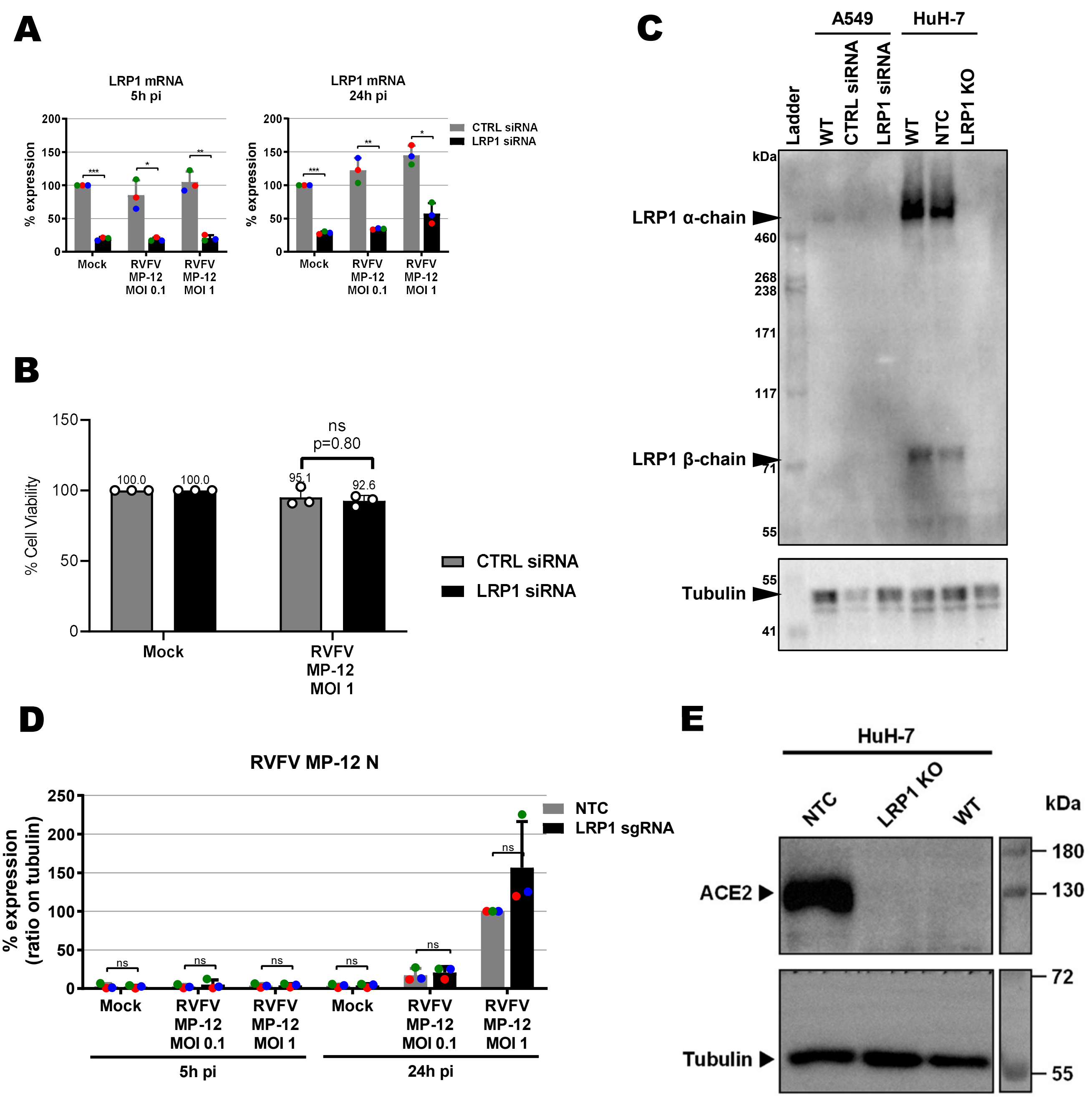


**Figure S5.** LRP1 depletion in A549 and HuH-7 cells. (A) LRP1 mRNA levels in control (CTRL) and LRP1 siRNA knock-down A549 cells as measured by RT-qPCR. Cells were infected with RVFV MP-12 at MOI 0.1 or MOI 1, total RNA was extracted at 5h and 24h post-infection (p.i.), and RT-qPCR was done to detect LRP1 and the GAPDH reference mRNAs. RNA levels in the CTRL mock-infected were set to 100%. (B) Cell survival. CTRL and LRP1 siRNA-treated A549 cells were infected with RVFV strain MP-12 at an MOI of 1, and cell survival measured 24 h later. (C) LRP1 protein levels in wild-type and knockdown or knockout A549 and HuH-7 cells, respectively, as measured by immunoblot analysis. NTC, no template control (clone E5); WT, wild-type. The shown LRP1 KO is from clone C8. (D) Quantification of RVFV N immunoblot signals from 3 independent experiments as shown in figure 3C. (E) ACE2 levels of the HuH-7 cells and cell clones shown in (C).

Figure S7

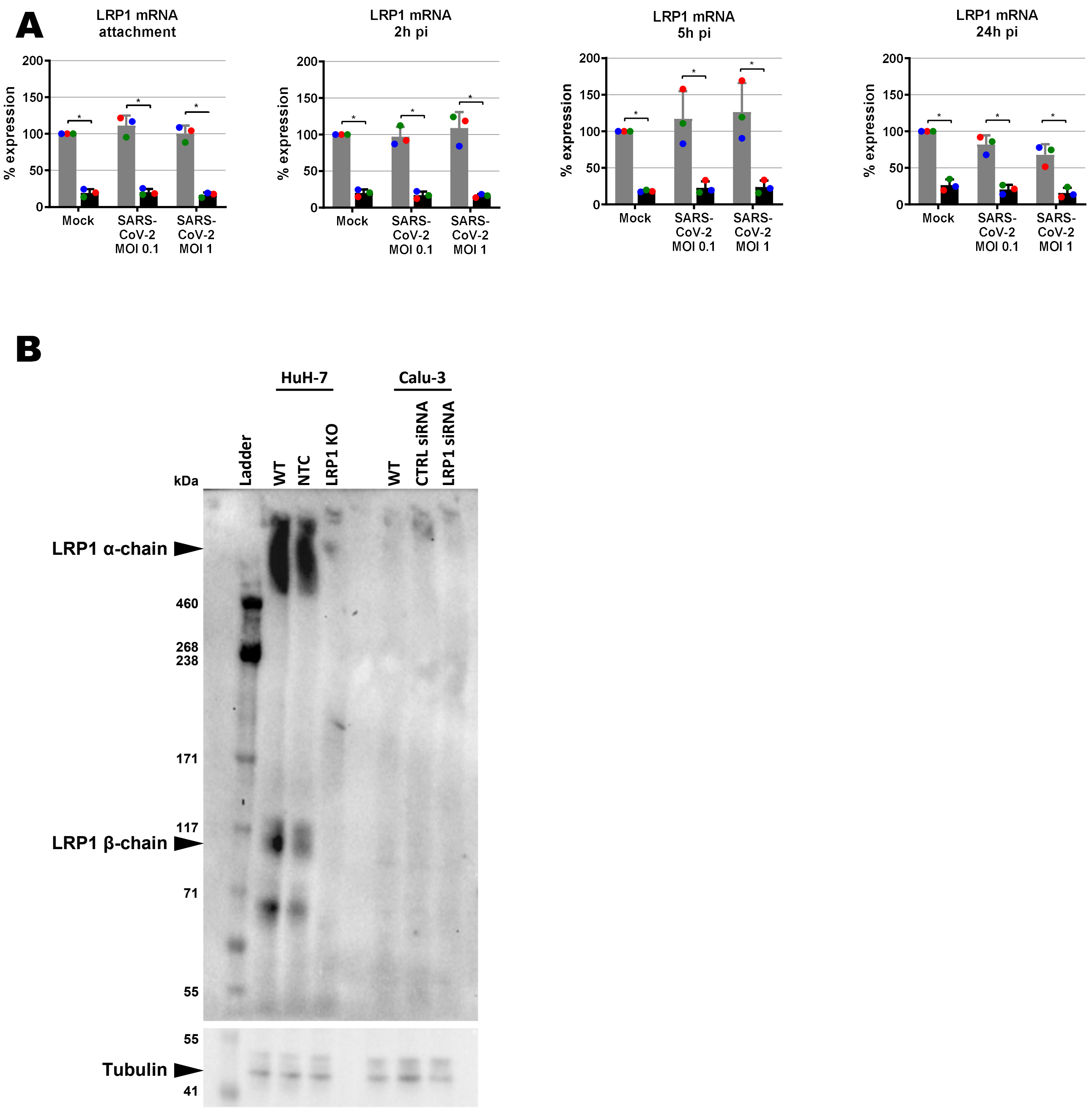


**Figure S6.** LRP1 expression in wt and LRP1-depleted human cell lines. (A) LRP1 mRNA levels in wild-type and knock-down Calu-3 cells as measured by RT-qPCR (samples of Fig. 6A). Cells were infected with SARS-CoV-2 at MOI 0.1 or MOI 1, RNA was extracted at the indicated time-points p.i., and RT-qPCR was performed to detect LRP1 and the GAPDH reference mRNAs. RNA levels in the CTRL mock-infected were set to 100%. (B) LRP1 protein levels in wild-type and knock-out HuH-7 cells and knock-down Calu-3 cells as measured by immunoblot analysis. CTRL, control; NTC, no template control; WT, wild-type.

Table S1

| Short name | Full name | Primer sequence | gDNA samples |
|---|---|---|---|
| **DS** | **iPCR_DS** | **AATGATACGGCGACCACCGAGATCTACACGAG CCAGAACCAGAAGGAACTTGAC** | **All** |
| **A1** | **iPCR_US_A_BC10** | **CAAGCAGAAGACGGCATACGAGAT*GTCGTC*GT GACTGGAGTTCAGACGTGTGCTCTTC** | **Surviving retro-library / MseI** |
| **T1** | **iPCR_US_T_BC10** | **CAAGCAGAAGACGGCATACGAGAT*AGTCGTC*G TGACTGGAGTTCAGACGTGTGCTCTTCC** | **Surviving retro-library / NlaIII** |
| **A2** | **iPCR_US_A_BC9** | **CAAGCAGAAGACGGCATACGAGAT*ACTACT*GT GACTGGAGTTCAGACGTGTGCTCTTC** | **Surviving wild-type / MseI** |
| **T2** | **iPCR_US_T_BC9** | **CAAGCAGAAGACGGCATACGAGAT*AACTACT*G TGACTGGAGTTCAGACGTGTGCTCTTC** | **Surviving wild-type / NlaIII** |

**Table S1.** Next Generation Sequencing (MiSeq, Solexa) iPCR primer sequences. Index sequence is marked in red italics.

Table S2

| sgRNA | sequence | location |
|---|---|---|
| **LRP1_fwd** | **CACCGGCATGGACGGCTCAGATGAG** | **Exon 3** |
| **LRP1_rev** | **AAACCTCATCTGAGCCGTCCATGCC** | |

**Table S2.** sgRNAs used for the CRISPR-hSpCas9 knock-out of the *LRP1* gene in HuH-7 cells.

### Table S3

| Gene | Primer/Probe | Sequence | Ref. |
|---|---|---|---|
| EMCV gene 2B | EMCV 2B-F | ATGGGAAAATGTAAAAGAAACA | (1) |
| | EMCV 2B-R | GCATCACTGCTATTGTCA | |
| | EMCV 2B-P | 6-FAM-AGCTGCACACATCTGCTCAA-BHQ1 | |
| LACV gene L | qPCR LACV fwd | AGGAAAACTCCTGAGAATATAACTA | This work |
| | qPCR LACV rev | GGTATACAAACTGGTGGCGAT | |
| | qPCR LACV probe | 6-FAM-CTTAAATTTGAAAATATGTCTAAAATCCAAACATACCCAGGC-BHQ-1 | |
| RVFV gene L | RVFL-2912fwdGG | TGAAAATTCCTGAGACACATGG | (2) |
| | RVFL-2981revAC | ACTTCCTTGCATCATCTGATG | |
| | RVFL-probe-2950 | 6-FAM-CAATGTAAGGGGCCTGTGTGGACTTGTG-BHQ1 | |
| SARS-CoV-2 gene E | E_Sarbeco_F | ACAGGTACGTTAATAGTTAATAGCGT | (3) |
| | E_Sarbeco_R | ATATTGCAGCAGTACGCACACA | |
| | E_Sarbeco_P1 | 6-FAM-ACACTAGCCATCCTTACTGCGCTTCG-BBQ | |
| SFSV gene L | SFT FP | TCTGAGAACTGAGCTACAAGTG TTTATTA | (4) |
| | SFT RP | TTCCCATCTCTCTTCTGAAGAGTG | |
| | SFT P | 6-FAM-AGGTCATAGACAGTATCATGAGAATTGCTAGGTG-BHQ-1 | |
| VSV gene N | VSV q-PCR fwd | GATAGTACCGGAGGATTGACGACTA | This work |
| | VSV q-PCR rev | TCAAACCATCCGAGCCATTC | |

**Table S3**. Primer and probe list for detection of virus RNA by qPCR.
